# Dysfunction of the Visual Sensory Thalamus in Developmental Dyslexia

**DOI:** 10.1101/2022.11.14.516174

**Authors:** Christa Müller-Axt, Louise Kauffmann, Cornelius Eichner, Katharina von Kriegstein

## Abstract

Developmental dyslexia (DD) is a reading disorder with a prevalence of 5-10%. Neuroscientific research has typically focused on explaining DD symptoms based on pathophysiological changes in the cerebral cortex. However, DD might also be associated with alterations in sensory thalami – central subcortical stations of sensory pathways. A post-mortem study on the visual sensory thalamus (lateral geniculate nucleus, LGN) showed histopathological changes in the magnocellular (M-LGN), but not in the parvocellular (P-LGN), subdivisions. M-LGN and P-LGN have different functional properties and belong to two different visual systems. Whether M-LGN alterations also exist in DD *in-vivo* is unclear. Also, the potential relevance of M-LGN alterations to DD symptoms is unknown. This lack of knowledge is partly due to considerable technical challenges in investigating LGN subdivisions non-invasively in humans. Here, we employed recent advances in high-field 7 Tesla functional magnetic resonance imaging (fMRI) to map the M- and P-LGN *in-vivo* in DD adults (n=26) and matched controls (n=28). We show that (i) M-LGN responses differ between DD and control participants, (ii) these differences are more pronounced in male than in female DD participants, and (iii) M-LGN alterations predict a core symptom of DD in male DD participants only, i.e., rapid naming ability. Our results provide a first functional interpretation of M-LGN changes in DD and support DD theories that propose a direct relevance of sensory thalamus alterations for DD symptoms. In addition, the sex-specific behavioral relevance of M-LGN alterations within DD calls for taking sex differences into account when planning brain-based therapeutic interventions.

**Significance Statement:** Developmental dyslexia (DD) is one of the most common learning disorders affecting millions of children and adults world-wide. Several decades ago, pioneering research in five DD post-mortem brains suggested that DD is characterized not only by alterations of the cerebral cortex, but also by changes in a subsection of the visual sensory thalamus – the so-called M-LGN. The relevance of these findings for DD remained highly controversial. Using recent developments in high-resolution functional neuroimaging, we now discovered that M-LGN alterations are present also in DD *in-vivo* and predict a core symptom of DD in males. Our results provide a first functional interpretation of M-LGN alterations in DD and provide a basis for better understanding sex-specific differences in DD.

## Introduction

Developmental dyslexia (DD) is a neurodevelopmental disorder characterized by persistent difficulties in acquiring effective literacy skills despite adequate intellectual development and educational opportunities (1). With a 5-10% prevalence in children, DD encompasses the most common learning disorder. DD is often associated with considerable long-term consequences for the individual and high costs for society (1, 2). Compared to typically reading peers, DD is associated with significantly higher academic drop-out and unemployment rates, poorer health, and a shortened life expectancy (1).

Research on the neurobiological origins of DD in humans focusses primarily on the cerebral cortex and has revealed alterations particularly in a left-lateralized language network (3). However, this cortico-centric view of DD is challenged by histopathological observations made in the early-1990s on several *post-mortem* brains of dyslexics (4, 5). These studies revealed that DD is not only associated with alterations (neuronal ectopias and focal microgyria) in key cortical language regions, but also with histological alterations of the sensory thalami, i.e., the lateral geniculate nucleus (LGN) and the medial geniculate body (MGB) of the visual and auditory processing pathway, respectively (4, 5). Sensory thalami are the last subcortical processing site before sensory information is routed to primary cortices (6). Sensory thalamus alterations were also observed in several animal models of DD (6, 7). *In-vivo* magnetic resonance imaging (MRI) studies in humans have shown predominantly left-hemispheric alterations of thalamo-cortical connectivity in DD in the visual and auditory pathway (9, 10).

In humans, the LGN is a small, layered structure that can be coarsely partitioned into two subdivisions: a magnocellular (M-LGN; layers 1-2) and a parvocellular (P-LGN; layers 3-6) subdivision (11, 12). Neurons of the two subdivisions process complimentary visual information: For example, M-LGN neurons are involved in coarse spatial image analysis and are specialized in detecting rapid visual changes and motion. Conversely, P-LGN neurons are involved in processing color and fine spatial detail (6). Human *post-mortem* studies in DD demonstrated morphological alterations specifically in the magnocellular layers but not in the parvocellular layers of the LGN (4). These findings were based on a relatively small number of *post-mortem* dyslexic cases (N=5) and have not yet been replicated in further human *post-mortem* or *in-vivo* imaging studies. Furthermore, not all dyslexics exhibit behavioral impairments that could be attributed to a general magnocellular visual processing difficulty (13). Thus, to date, it remains elusive (i) whether M-LGN alterations can also be detected in DD *in-vivo*, and if so, (ii) which functional relevance these may have for dyslexia symptoms.

The scarcity of clinical *post-mortem* brain specimens and the technical challenges associated with *in-vivo* MRI measurements of small subcortical brain structures pose major obstacles to answering these questions. In humans, individual LGN layers are ≤1mm thick, verging on the limits of attainable image resolutions of conventional MRI (11, 12). However, recent advances in high-field MRI have made it possible to measure distinct signals from the M- and P-LGN in humans *in-vivo*, paving the way for assessing subdivision-specific LGN alterations in larger sample sizes (12, 14).

Using recently developed high-field functional MRI (fMRI) experiments at 7 Tesla, we investigated whether DD is associated with functional alterations of the M-LGN. We acquired data from a large sample of N=54 young German adults with a lifelong history of DD and matched control participants (Table S1). With this sample we performed three 7T blood oxygen level-dependent (BOLD) fMRI experiments (Fig. 1). The central aim of these experiments was to test whether (i) M-LGN alterations can also be detected in DD *in-vivo*, and if so, (ii) whether they are related to a dyslexia diagnostic score, i.e., rapid automatized naming for letters and numbers (RANln). RANln performance is key for predicting reading ability (15) and is associated with alterations of connections between LGN and cerebral cortex in DD (10) as well as thalamo-cortical alterations in the auditory modality (16).

**Figure 1.**
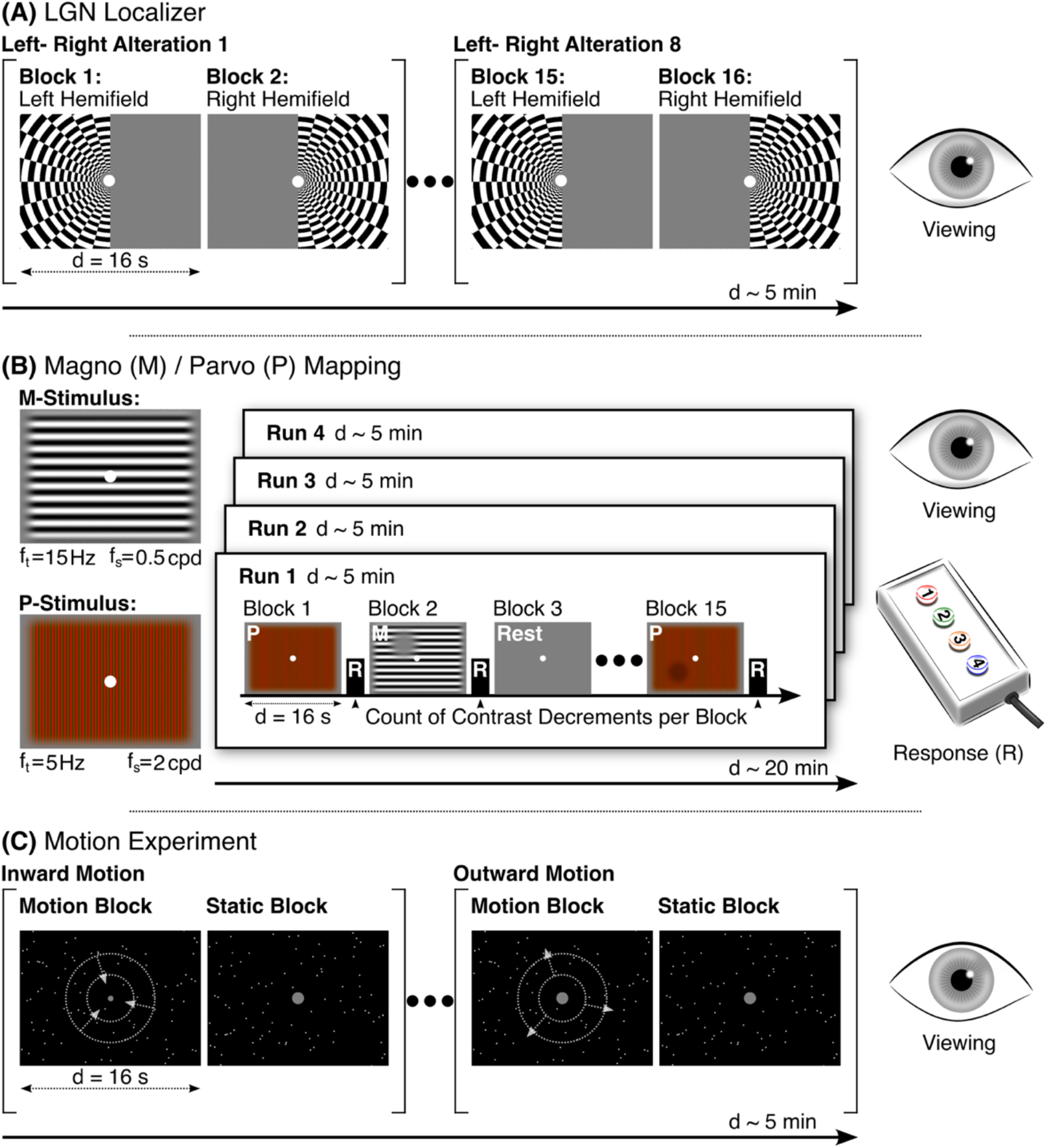
Experimental design of the three fMRI experiments. **(a)** In the LGN localizer, participants saw a flickering checkboard stimulus in blocks alternating between the left and right visual hemifields. They viewed the stimuli while maintaining central fixation. **(b)** During the M/P mapping experiment, participants viewed two types of experimental stimulus blocks which were designed for evoking different BOLD responses from the M- and P-LGN. M-blocks consisted of a full-field achromatic grating stimulus, presented at low spatial (f_s_) and high temporal frequency (f_t_). P-blocks consisted of a full-field colored grating stimulus, presented at higher spatial (f_s_) and lower temporal frequency (f_t_). M- and P-blocks were interleaved with rest blocks containing a gray screen. During the experimental stimulus blocks, participants had to detect contrast decrements and report the number of targets within a block (luminance in M blocks, color in P blocks) by button press after each block (R). **(c)** In the visual motion experiment, participants saw blocks of either moving or static point clouds. Blocks with moving point clouds consisted of either inward or outward motion. Participants viewed the stimuli while maintaining central fixation. See Materials & Methods for more details on the experimental designs. Abbreviations: LGN, lateral geniculate nucleus; d, duration; M, magno; P, parvo.

## Results

In each individual participant, we segmented the entire LGN based on an anatomical atlas and additionally localized it functionally (LGN localizer, 12) (Fig. 1A). Within the LGN, we mapped the M- and P-LGN (M/P mapping, 12) (Fig. 1B) to test for functional differences in the M- and P-LGN between control and DD participants. In a further fMRI experiment, we assessed visual motion processing (visual motion experiment) (Fig. 1C). This experiment was originally developed in the context of a different research question as a V5/MT-localizer and here served as a quality control for the identified M- and P-LGN derived from the M/P mapping experiment.

### Overall LGN Responses Similar Between Control and DD Participants

The LGN localizer (Fig. 1A) allowed us to functionally localize the entire LGN in each participant and to assess whether DD participants may already differ from control participants in their overall functional LGN responses to visual stimulation. Such a difference between groups would indicate a general LGN deficit in DD that is not confined to any particular LGN subdivision. A mixed-design analysis of variance (ANOVA) of participants’ functional LGN responses with the between-subject factor of group (controls/DD) and the within-subject factors of hemisphere (left/right) and stimulation site (left/right visual hemifield) provided no support for such a global LGN deficit in DD. There was neither a significant main effect (*F*(1,52) = 0.132, *p* = .718, 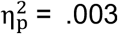) nor any interaction (all *p*’s ≥ .200, all 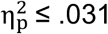) with the factor group, suggesting that overall LGN responses to the visual stimulation were similar in control and DD participants.

### Altered M-LGN Response in DD Participants

We then addressed whether DD is associated with specific alterations of the M-LGN. For this purpose, we functionally defined each participant’s M- and P-subdivision (Fig. 2A/B). We then computed a mixed-design ANOVA of participants’ subdivision-specific LGN responses with the between-subject factor of group (controls/DD) and the within-subject factors of subdivision (M-LGN/P-LGN), stimulus-type (M-stimulus/P-stimulus), and hemisphere’(left/right). The analysis revealed a significant three-way interaction of group × subdivision × hemisphere (*F*(1,47) = 4.974, *p* = .031, 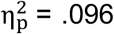; Fig. 3). The observed three-way interaction suggested a difference in the subdivision × hemisphere interaction between the two groups. Given previous results of potential left lateralization of sensory thalamus alterations in DD (9, 10, 16), we expected a significant subdivision × hemisphere interaction in the DD but not in the control group. In line with this expectation, two subsequent within-group repeated-measures ANOVAs revealed a significant interaction of subdivision × hemisphere in DD participants (*F*(1,24) = 6.531, p = .017, 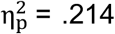, at Bonferroni-adjusted significance level a = .025), while this interaction was non-significant in controls (*F*(1,23) = 0.724, *p* = .403, 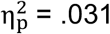) (Fig. 3). Post-hoc paired t-tests showed that the observed subdivision × hemisphere interaction in the DD group was driven by significant hemispheric differences in functional responses between the left and right M-LGN (*t*(24) = 3.199, *p* = .004, *d* = 0.64, two-tailed with Bonferroni-adjusted significance level a = .025), but not the P-LGN (*t*(24) = 0.520, *p* = .608, *d* = 0.104) (Fig. 3). M-LGN responses in DD participants were significantly stronger in the left than right hemisphere. Overall, these results suggest that unlike typical readers, DD participants have functional response alterations that specifically affect the M-LGN. Consistent with earlier *post-mortem* human studies (4), these findings provide first evidence that DD is associated with alterations of M-LGN also *in-vivo*.

**Figure 2.**
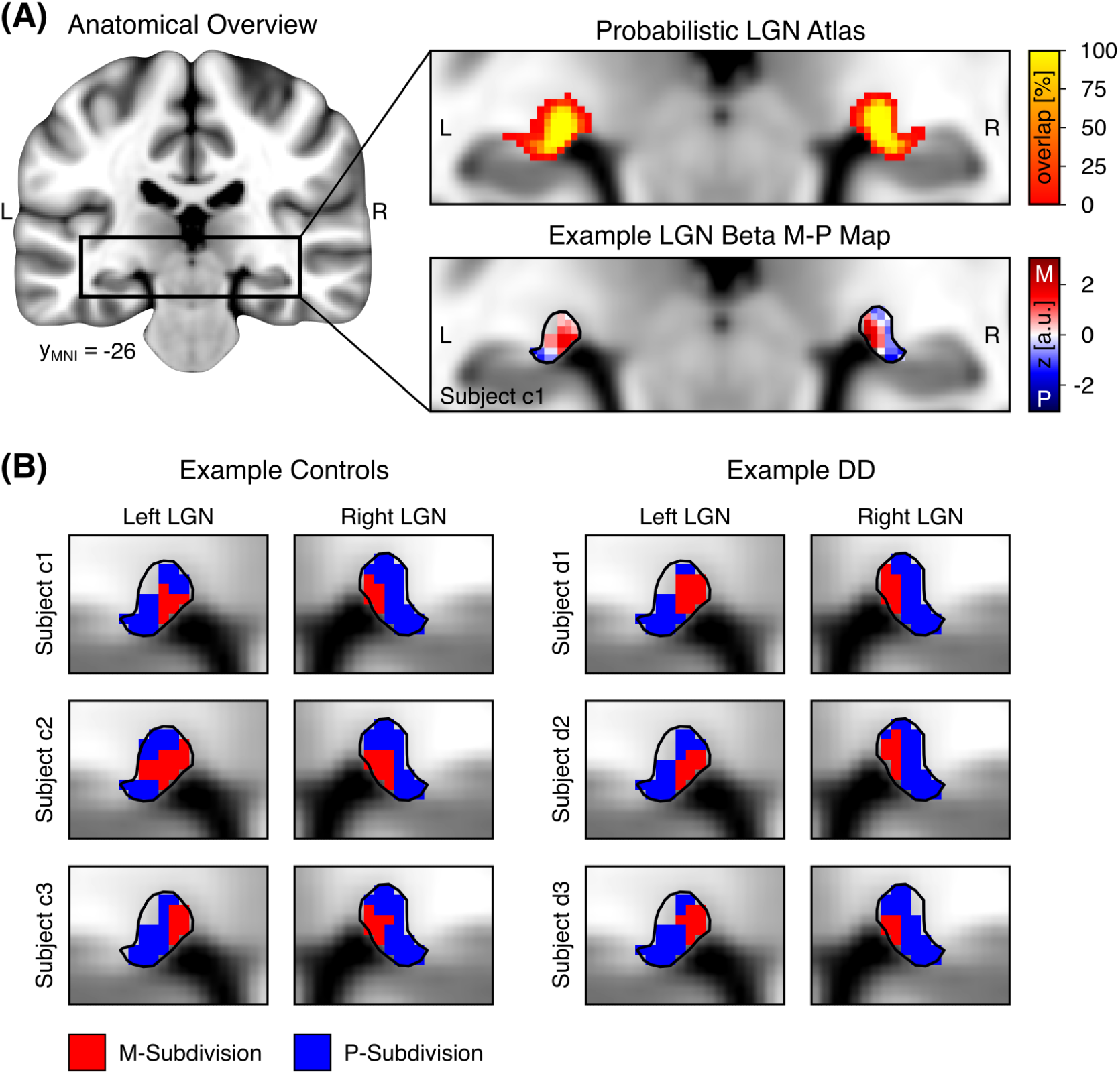
Definition of M/P-LGN in control and DD participants. **(A)** Left: Anatomical overview of the location of the LGN, indicated by the black rectangle, within the standard brain of the Montreal Neurological Institute (MNI). Right: We used a publicly available high-resolution probabilistic LGN atlas (top panel) to confine functional responses from the LGN localizer to the bilateral nuclei in each participant (1). The atlas was set to a threshold of 35% overlap across subjects, indicated by the solid black outline around the LGNs (bottom panel). Within these defined regions, M/P-LGN mapping was performed by 20/80% volume thresholding of the obtained BetaM-P maps (bottom panel) from the M/P mapping experiment (2). On the BetaM-P map, LGN voxels with larger values (red color) show a higher response preference for the M-stimulus, while voxels with lower values (blue color) show a higher response preference for the P-stimulus. **(B)** Examples of derived M-LGN (in red color) and P-LGN maps (in blue color) based on volume thresholding in individual representative control and DD participants. Abbreviations: a.u., arbitrary units; LGN, lateral geniculate nucleus; M, magno; P, parvo; L, left; R, right.

**Figure 3.**
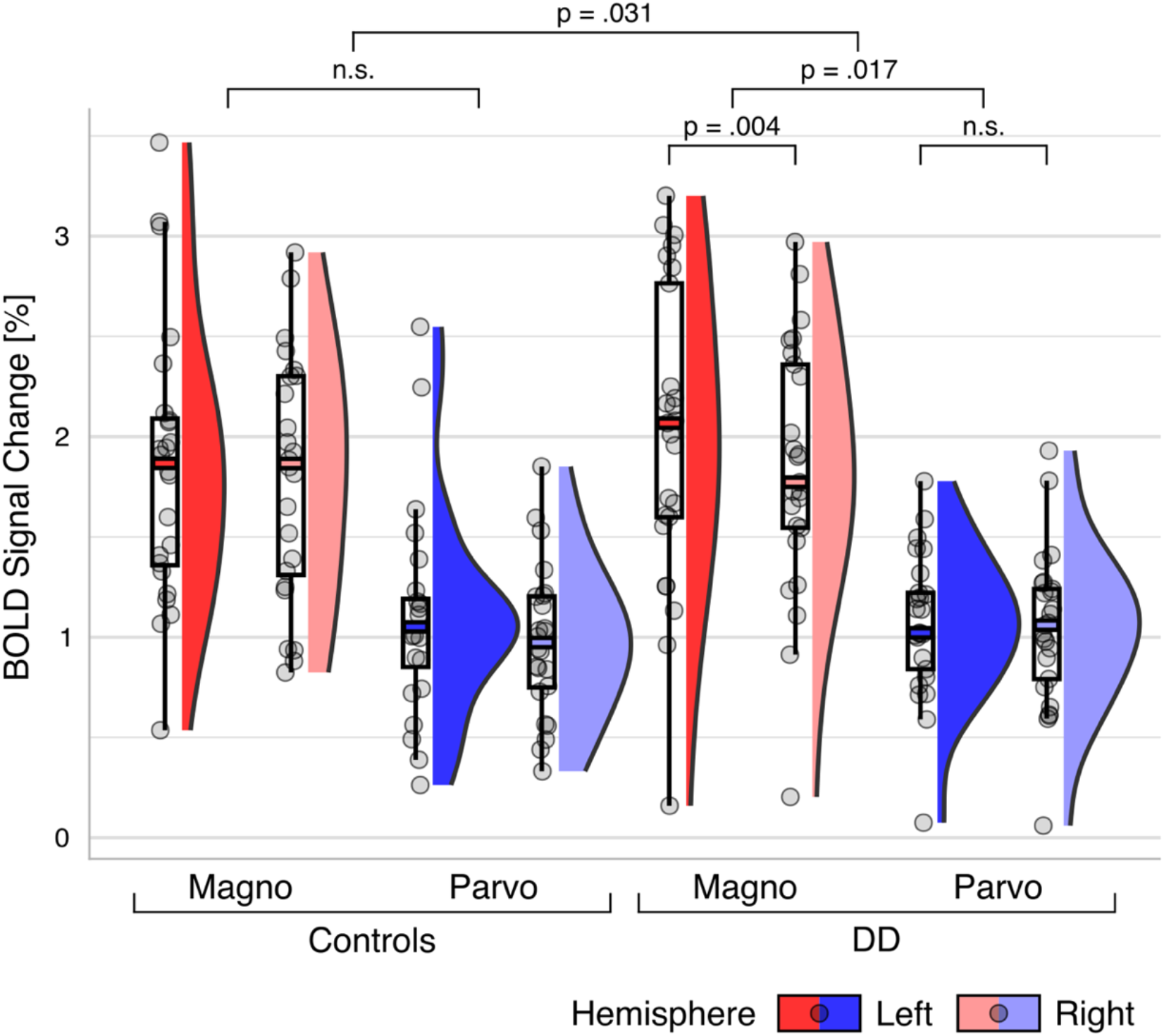
Bilateral M/P-LGN BOLD responses in control (n=24) and DD (n = 25) participants. The figure displays boxplots overlaid with individual data points alongside color-coded density plots. M-LGN responses are coded in red. P-LGN responses are coded in blue. Left and right-hemispheric responses are coded in dark and light color, respectively. Responses are averaged across stimulus-type to reveal the significant interaction between group x subdivision x hemisphere. Abbreviations: n.s., not significant; BOLD, blood oxygen level-dependent.

### No Differences in M/P-LGN Localization Accuracy Between Groups

Our finding of stronger left than right-hemispheric M-LGN BOLD responses in DD but not in control participants cannot be explained by group differences in the localization strategy or accuracy of the LGN masks (Fig. 2A) or M/P-LGN maps (Fig. 2B). First, there were no significant differences between control and DD participants in the size of the individually defined entire LGN masks (indicated by solid black LGN outlines in Fig. 2), neither for the left LGN (128.2 ±17.4 mm^3^ in controls vs. 124.5 ±14.0 mm^3^ in DD; *t*(52) = 0.838, *p* = .406, *d* = 0.228, two-tailed) nor for the right LGN (136.1 ±17.0 mm^3^ in controls vs. 132.4 ±17.7 mm^3^ in DD; *t*(52) = 0.793, *p* = .432, *d* = 0.216, two-tailed). Second, all M/P-LGN maps were subjected to the anatomically informed criterion that, for each participant, the identified M-LGN should be consistently located more medial than the identified P-LGN (12, 14) (Fig. 2B) (SI Methods). Participants for whom this was not the case (n = 4 controls, n = 1 DD) were excluded from the above analysis. There were no differences in the size of the identified M/P-LGN maps between groups in either hemisphere in the final sample (all *p*’s ≥ .512, all *d’s* ≤ 0.189). Also, the behavioral performance on the contrast decrement detection task during the M/P mapping experiment did not significantly differ between groups (all *p*’s >0.4; SI Results).

Lastly, we verified that the identified M/P-LGN maps also showed the expected functional response properties of M-LGN and P-LGN (SI Methods). To this end, we analyzed the subdivision-specific LGN responses to an independent visual motion stimulus. Based on the known response properties of M-LGN and P-LGN neurons, BOLD responses to visual motion should be stronger in the identified M-than P-subdivisions (6). As expected, a mixed-design ANOVA of participants’ functional M/P-LGN responses to the contrast motion vs. static with the between-subject factor of group (controls/DD) and the within-subject factors of subdivision (M-LGN/P-LGN) and hemisphere (left/right) revealed a significant main effect of the factor subdivision with stronger BOLD responses in the identified M-than P-subdivisions across participants (*F*(1,46) = 57.621, *p* = 1.194×10^-9^, 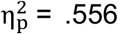). There was no main effect (*F*(1,46) = 0.181, *p* = .673, 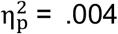) nor any interaction (all *p*’s ≥ .268, all 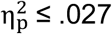) with the factor group, suggesting that the identified subdivisions in both groups adhered to the expected functional response properties.

### Sex Differences in M-LGN Response in DD

Previous human *post-mortem* and *in-vivo* MRI studies demonstrating sensory thalamus alterations in DD have been based almost exclusively on all-male DD cohorts (4, 5, 9, 10, 16). This aspect is intriguing, as several findings from animal models suggest that there may be hormone-related differences in the extent of sensory thalamic alterations between the sexes in DD (7, 8, 17, 18). In particular, thalamic alterations in animal models of DD are related to gestational testosterone levels and, consequently, are more likely to affect male than female individuals (17).

Next, we thus explored whether the M-LGN alterations, quantified as a difference score between the functional BOLD responses of the left and right M-LGN (“M-LGN difference score”), differed between male and female participants in our DD sample. In line with the findings from DD animal models, an independent t-test revealed that M-LGN difference scores were indeed significantly larger among male than female DD participants (*t*(22) = 2.522, *p* = .019, *d* = 1.033, two-tailed) (Fig. 4A).

**Figure 4.**
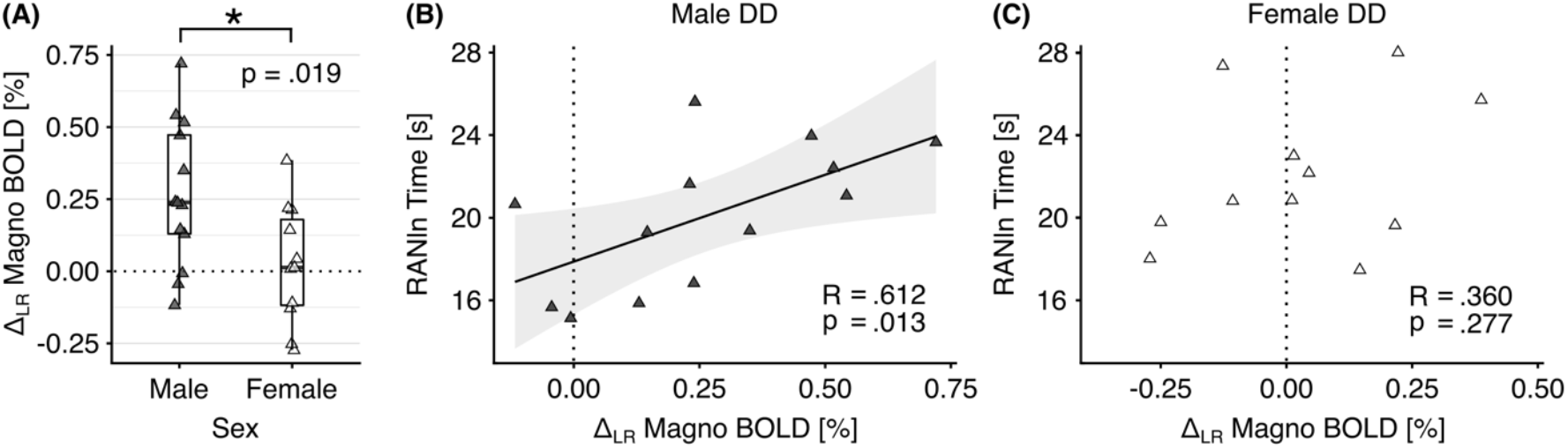
M-LGN response in DD participants (N = 24), and its behavioral relevance for rapid automatized naming for letters and numbers (RANln). **(A)** M-LGN response, quantified as a difference score between the BOLD responses of the left and right M-LGN (i.e., Δ_LR_ Magno BOLD) in male (n = 13, dark triangles) and female participants (n = 11, bright triangles) with DD. The dotted line indicates equal functional contributions of the left and right M-LGN to the difference score (i.e., no functional lateralization). **(B,C)** M-LGN difference scores correlate positively with the reaction time on RANln in male DD (A), but not in female DD participants (C). The plot in (B) shows the least squares correlation fit, including the 95% confidence interval (light gray shaded area) for the correlation coefficient R, in male participants with DD. Abbreviations: BOLD, blood oxygen leveldependent.

### M-LGN Response Predicts Key Deficit in DD Males

In human DD research, a commonly used diagnostic task is the RAN-task. In this task, participants name a series of visually presented familiar items (e.g., letters and numbers) aloud as quickly and accurately as possible (19). RAN ability is an important predictor of reading fluency and poses a key deficit in DD across the lifespan (15). Importantly, slow reaction times on RAN for letters and numbers (RANln) have previously been linked to both functional and structural alterations of the sensory thalami and their connections to cortex in DD (10, 16). We therefore expected that the reaction times on RANln would be associated with M-LGN alterations in DD. In this context, an interesting aspect discovered in animal models of DD is that only those animals that exhibited thalamic alterations also showed behavioral impairments (7, 17). Furthermore, previous studies on the association between RANln and thalamo-cortical alterations in DD relied predominantly on male samples, limiting their predictive power for similar associations in female DD. We therefore correlated the M-LGN difference scores with the reaction times on RANln using one-tailed Pearson’s correlations across the whole DD group, and within male and female DD participants separately. The analyses revealed, in male DD participants only, a significant correlation between M-LGN difference scores and RANln performance (*R* = .612, *p* = .013, at Bonferroni-adjusted significance level a = .0167) (Fig. 4B). The correlations across the whole DD group (*R* = .293, *p* = .082) and within female DD participants (*R* = .36, *p* = .277) were non-significant (Fig. 4C).

## Discussion

Recent developments in high-field MRI have enabled the study of small brain structures such as the subdivisions of human thalamic nuclei *in-vivo*. We here used this technical advance to image the human LGN and its M and P-subdivisions in a large sample of adults with DD and matched control participants. Consistent with human *post-mortem* reports dating back to the 1990s (4, 5), we found that individuals with DD show functional response alterations specifically in the M-LGN. Our findings solve the long-standing question whether M-LGN alterations are also present in DD *in-vivo* and give first indications about their behavioral relevance as well as sex-dependency.

Our finding of different lateralization of the M-LGN in DD in comparison to controls parallels previous findings of left-lateralized sensory thalamic alterations in DD (5, 9, 16). In the auditory pathway, histological changes occurred specifically in the left MGB in *post-mortem* brains of dyslexics (5). Also, *in-vivo* MRI studies on DD showed functional response changes and altered connectivity of the MGB restricted to the left hemisphere (9, 16). Previous findings on potential laterality of thalamic alterations in the visual processing pathway are less conclusive (4, 10): histopathological changes were found in the M but not in the P layers of the LGN; however, it is unclear which hemisphere(s) were affected (4). In addition, there is reduced structural connectivity between the left LGN and visual motion area V5/MT in DD, however connectivity results in the right hemisphere remained unclear (10). Recent behavioral findings point towards an altered lateralization in DD also in visual processing: while typically reading individuals have a right hemifield advantage in detecting moving low-spatial frequency events, this is not the case in DD (20). Our results do not permit to adjudicate whether the divergent lateralization of the M-LGN is due to response differences within the left or the right M-LGN. However, given the left-lateralized auditory thalamic changes *in-vivo* and *post-mortem* (5, 9, 16), the aberrant left-hemispheric cortico-thalamic LGN-V5/MT connectivity (10), and first indications from behavioral findings (20), we suggest that thalamic changes in the visual processing pathway in DD may be primarily left-lateralized.

Animal models of DD have shown sex differences in the extent of thalamic alterations and their relation to behavioral impairments: due to higher testosterone levels during gestation, male animals are more likely affected by sensory thalamic alterations and associated behavioral deficits than female animals (7, 17). Our findings are the first indication that similar sex differences might also occur for thalamic alterations in human DD. We found that functional responses of the M-LGN related to a key deficit in DD (i.e., RANln), particularly in males. Impaired RANln performance has been repeatedly associated with left-hemispheric sensory thalamic alterations in DD in previous studies (10, 16). These studies were, in fact, consistently based on all-male DD cohorts. The correlation between RANln and sensory thalamus alterations observed in our and previous studies may be a hallmark of DD that is predominant in male individuals. Sensory thalamus alterations may contribute to the higher prevalence of DD in males than in females (ratio 3:1) (3). These findings stress the need for more sex-specific brain models of DD in a research area otherwise heavily skewed toward males (21–23).

We cannot derive from our results *how* thalamic alterations contribute to core DD symptoms. We have previously suggested two possible explanations (10). First, successful reading and RANln performance involve rapid attentional shifts toward successive visual-spatial cues – a skill largely controlled by a right-lateralized frontoparietal attention network (24, 25). Neurons of the M-LGN relay visual information via the dorsal stream to area V5/MT, which in turn serves a major input structure to this attention network (26). The association between the left-lateralized M-LGN responses and RANln performance in DD could thus result from deficient attentional mechanisms (27, 28) through inefficient interactions with this typically right-lateralized attention network. Our second suggestion was that deficient RANln performance might be a result of deficient top-down modulation of the LGN to fast-varying predictable speech stimuli, i.e., visual articulatory movements (10). M-LGN neurons are known to process high temporal frequency visual information (6). Interestingly, DD is associated with a reduced structural connectivity between the LGN and area V5/MT in the left hemisphere (10). An imbalanced top-down modulation of LGN-M neurons could therefore contribute to a deficit in processing fast visual speech features in DD, which might be important for acquiring phonologic skills during ontogeny.

In summary, our results show that M-LGN alterations are a key feature of DD and are associated with reading-related behavioral scores, particularly in male DD. The findings suggest that (i) sex differences in the brain basis of DD extend beyond the cerebral cortex to the sensory thalamus, and (ii) that an understanding of sensory thalamus alterations in DD would benefit from a thorough understanding of sex-related developmental determinants of thalamic maturation. The findings are also relevant for clinical studies as they suggest that targeting the thalamo-cortical system for example with complementary neurostimulation might be particularly effective in male individuals with DD (29, 30).

## Materials and Methods

### Participants

Fifty-four healthy adult German speakers were included in the analyses. This sample consisted of 26 participants with DD and 28 control participants, matched in age, sex, handedness, and nonverbal IQ (SI Methods and Table S1). Participants with DD performed worse than controls on tests of literacy (spelling, reading speed and comprehension), rapid automatized naming of letters and numbers (RANln), and word and non-word reading (SI Methods and Table S1).

### MRI Experiments: Procedure

Participants attended two MRI sessions on two separate days. The sessions included three fMRI experiments: the LGN localizer and M/P mapping experiment (first session) and a motion experiment (second session). In addition, a set of whole-brain quantitative structural MR images were acquired in each participant during the first session. One DD participant attended only the first MRI session due to pregnancy at the time of the second session. In the context of a different research question, we acquired additional fMRI and diffusion-weighted imaging data from the participants, the results of which will be reported elsewhere.

### MRI Experiments: Setup

In each fMRI experiment, visual stimuli were front-projected onto a translucent screen positioned on the participants’ chest. Participants viewed the screen in the MRI system through a mirror mounted just above their eyes. During the fMRI experiments, we also recorded cardio-respiratory data from the participants. This was done to account for physiological noise in the BOLD signal during data processing in order to increase the signal-to-noise ratio in the LGN (31). For more details on display settings, visual stimulation software, and physiological recordings, see SI Methods.

### Experimental Design: LGN Localizer

This experiment was used to functionally localize the LGN in each participant (14) (Fig. 1A). The stimulus consisted of a flickering radial checkerboard with 100% contrast, with its contrast polarity reversed at 4Hz (for the full cycle). The checkerboard covered half the screen while the other half contained a uniform gray background. The checkerboard alternated between the two visual hemifields in a block-design fashion. Participants maintained fixation on a central white fixation dot while viewing the stimuli. Each hemifield block lasted 16 s and the whole run was composed of 8 left-right alternations for a total of 16 blocks and a run duration of 5 mins. Further details on the experimental design can be found in the corresponding reference (14).

### Experimental Design: M/P Mapping

This experiment used full-field stimuli designed to match the selective response properties of neurons in the M-LGN and P-LGN (14) (Fig. 1B). The M-stimulus was a sinusoidal grayscale grating with a luminance contrast of 100%, a low spatial frequency of 0.5 cpd, and a sinusoidal counterphase flicker frequency of 15 Hz. The P-stimulus was a sinusoidal high color-contrast red-green grating with low luminance contrast, a higher spatial frequency of 2 cpd, and a lower sinusoidal counterphase flicker of 5 Hz. Gratings changed orientation every 3 s and could be presented at one of 6 orientations (0°, 30°, 60°, 90°, 120°, 150°). M and P-stimuli were presented in a blocked design and were interspersed with rest blocks consisting of a uniform gray background. Throughout the experiment, participants maintained fixation on a central white fixation dot while viewing the stimuli. To ensure continued fixation on the screen during experimental M/P-blocks, participants were asked to detect contrast decrements (0 to 3 targets) that could appear at random locations within each block. At the end of each block, participants had 1.5 s to report the number of targets per button press. Each block lasted 16 s and each run was composed of 6 M blocks, 6 P blocks and 3 rest blocks for a total of 15 blocks. Participants completed 4 runs of the M/P mapping experiment, which lasted approximately 5 mins each. Further details on the experimental design can be found in the corresponding reference (14).

### Experimental Design: Motion Experiment

This experiment served to functionally validate the obtained M- and P-subdivision maps and consisted of alternating moving and static point clouds presented in a block design (Fig. 1C). In the motion blocks, point clouds consisted of 250 white dots with a radius of 0.1° moving radially against a black background at a speed of 4.7 deg/s and 100% coherence within a circular aperture of 17°. For half of the motion blocks, the points moved inward, while for the other half, they moved outward. Radial motion was chosen to facilitate central fixation and to stimulate a broad spectrum of motion direction-selective cells (32). During static blocks, the same number of dots were displayed at random locations and remained stationary over the duration of the block. Throughout all blocks, participants were instructed to maintain fixation on a central gray fixation point (0.2° of radius) while viewing the stimuli. Each block lasted 16 s and a run was composed of 8 blocks of each type (i.e., motion and static) for a total of 16 blocks. Participants completed one run which lasted ~5 mins.

### High-Resolution MRI Data Acquisition

High-resolution functional and structural MRI data were acquired on a 7 Tesla Magnetom MRI system (Siemens Healthineers, Erlangen, Germany) equipped with a 32-channel head coil (Nova Medical, Wilmington, MA, USA). In the three fMRI experiments, high-resolution echo-planar images were acquired at a resolution of 1.25 x 1.25 x 1.2 mm with partial brain coverage (40 transverse slices) covering the LGN and visual cortex. High-resolution whole-brain quantitative structural MR images were acquired (0.7 mm isotropic resolution) for registration purposes and as anatomical reference. Participants received foam padding around the head to reduce head motion. For further details on the acquisition parameters and a quantitative evaluation of head motion, see SI Methods.

### MRI Data Processing

Preprocessing and 1st-level statistical analyses of fMRI data were performed using standard pipelines in SPM12 (Wellcome Centre for Human Neuroimaging, London, UK), implemented in Matlab 2019Rb (Mathworks Inc., Sherborn, MA, USA) (SI Methods).

### Definition of the LGN

We used a publicly available, high-resolution 7T probabilistic LGN atlas (12) to precisely segment the LGN in each individual participant and to carefully demarcate it from adjacent visual brain structures. Nonlinear registrations of the atlas to each participant’s native quantitative T_1_ image were performed using (landmark-based) symmetric normalization in ANTs (version 2.3.1, 33; SI Methods). For each participant, individual left and right LGN masks were then registered to the functional image data. We also verified whether the resulting masks overlapped with the functional responses obtained in the LGN localizer experiment.

### Definition of M- and P-LGN

M- and P-LGN were defined using the M/P mapping experiment as previously described (14): For each participant, we computed Beta M-P maps in native space by subtracting the Beta maps obtained from the general linear model (GLM) estimation corresponding to the M- and P-stimulus conditions of the M/P mapping experiment, respectively. It follows that voxels with larger values on the Beta M-P maps correspond to a higher response preference for the M-stimulus, while voxels with lower values correspond to a higher response preference for the P-stimulus. To confine these maps to relevant voxels within the LGN, individual Beta M-P maps were then masked with the previously defined individual left and right LGN masks. For each participant and hemisphere separately, the M-LGN was defined as the 20% of voxels with the largest Beta M-P values, while the remaining 80% of voxels formed the P-LGN. This 20/80% volume allocation criterion is based on previous histological studies showing that the proportion of M and P neurons in the human LGN fall within these bounds, respectively (11).

As a quality criterion, we checked whether the M/P subdivision maps defined in native space adhered to the anatomically known spatial configuration of the M-LGN being located more medially than the P-LGN (12, 14). To do this, we computed individual M/P subdivision maps also in MNI standard space. This step permitted comparability between participants by aligning all LGNs in a common reference space. MNI Beta M-P maps were masked by a probabilistic LGN atlas (12, in MNI 1mm standard space) (Fig. 2A) and the same 20/80% volume allocation criterion was applied to define M- and P-LGN maps, respectively (Fig. 2B) (SI Methods). The MNI standard space analysis only subserved the quality control analysis of the spatial configuration of LGN-M/P maps. All reported quantitative analyses on the M/P-LGN localization accuracy between groups are based on the data in participants’ native space.

### Extraction of signal change

Beta estimates corresponding to the conditions of interest were extracted from all voxels within the LGN (i.e., for the LGN localizer experiment) or the M- and P-LGN (i.e., for the M/P mapping and motion experiments) in each participant using an in-house toolbox and converted to % signal change. The % signal change (PSC) was computed as: PSC = β_condition_ x SF / β_constant_ x 100 (wherein: β_condition_ = parameter estimate of the condition of interest, β_constant_ = parameter estimate for the constant term, SF = scale factor of the design matrix) (34). Finally, the mean PSC within each region was extracted for each participant and experimental condition of the fMRI experiments and subjected to mixed-design ANOVAs for statistical analysis (SI Methods). For the motion experiment, we used the M- and P-LGN to mask responses in the contrast motion vs. static.

## Data Availability

The scripts used to generate the LGN hemifield and M/P stimuli are publicly available (14). The motion experiment and fMRI analysis scripts have been made publicly available on the Open Science Framework (osf; https://osf.io/bge75/). Raw MRI data cannot be made available as sharing these personal data is not covered by the ethics clearance. Single-subject data in native and MNI space (i.e., individual LGN, M- and P- as well as beta M-P maps) are available on osf.

## Acknowledgments and Funding Sources

We are grateful to our participants for taking part in this study. We also thank the German federal association for dyslexia and dyscalculia (Bundesverband Legasthenie Dyskalkulie e.V., https://www.bvl-legasthenie.de/) for their continued support in advertising our study to potential participants. We further thank Martina Dietrich, Liane Dörr, Laura Hüser, Kim Lawatsch, Lisa Ziggel, Hanna Ovesiek, and Tabea Fleps for their extensive help in scheduling and conducting the behavioral tests. This work was supported by the ERC-consolidator-grant SENSOCOM 647051.

## Supporting Information

### SI Methods

#### Participants

All participants were tested on literacy skills, including reading speed and comprehension (LGVT; 1) and spelling (RT; 2), as well as on rapid automatized naming of letters and numbers (RANln; 3), and word and non-word reading (4) (Table S1). Participants provided written informed consent before study participation. The study was approved by the ethics committee of the Medical Faculty, University of Leipzig, Germany.

#### Inclusion Criteria for DD Participants

Participants with DD were required to meet the following criteria to be included in the study: (i) reading accuracy and/or speed, as assessed by measures commonly used for diagnosis of DD in Germany (i.e., LGVT or non-word reading), of at least 1.5 standard deviations (SD) below the mean of the matched control group; and (2) a life-long history of DD in the anamnesis. Participants with DD were recruited nationwide through print and online study advertisements.

#### General Participant Inclusion Criteria

All participants had to fulfill the following inclusion criteria: (i) no prior history of neurological and/or psychiatric disorders, (ii) free of psychostimulant medication, (iii) no co-existing neurodevelopmental disorders other than dyslexia (e.g., dyscalculia, autism spectrum disorder), (iv) no hearing disabilities, (v) normal or corrected-to-normal visual acuity, and (vi) a non-verbal IQ ≥ 85.

The first four criteria were assessed based on participants’ self-reports and screening questionnaires including the Autism-Spectrum Quotient (AQ; 5) and a brief, self-designed 10-item questionnaire on the main symptoms of dyscalculia (6, 7). Visual acuity was assessed through the Freiburg Visual Acuity Test (FrACT3; 8, 9; https://michaelbach.de/fract/) with a cutoff of +0.1 binocular logMAR to ensure normal visual acuity (10). Non-verbal IQ was assessed with the German adaptation of the Wechsler Adult Intelligence Scale-Revised (HAWIE-R; 11).

Finally, all participants had to meet the local safety requirements for high-field MRI: no metal implants, free of tattoos and non-removable ferromagnetic jewelry, no dental amalgam restorations, complete medical documentation of all potentially relevant previous surgical procedures and accidents, and no pregnancy in female participants (with the option to perform a rapid pregnancy test on site).

#### Display and Visual Stimulation Software

Within the MRI system, participants viewed the screen from a total viewing distance of 35 cm, which subtended approximately 18 x 16 degrees of visual angle. Stimuli were generated on Linux using the Psychtoolbox (12, 13), implemented in GNU Octave, version 4.2.0 (14), and presented at a refresh rate of 60 Hz.

#### Physiological Data Recordings

We recorded participants’ cardio-respiratory data throughout each fMRI experiment using an MRI-compatible Biopac System (Biopac Systems, Inc., Goleta, CA, USA). Cardiac signals were recorded through a pulse oximeter placed on participants’ left index finger with a sampling frequency of 100Hz. Respiratory data were recorded through thoracic movements using a non-electrical pressure pad placed on participants’ chest in combination with a respiration transducer. MR trigger pulses were also recorded to synchronize physiological parameters to each MR volume.

#### High-Resolution 7 Tesla Functional MRI Acquisition

High-resolution functional MRI data were acquired using a gradient-echo echo-planar imaging (EPI) sequence with the following imaging parameters: 1.25 mm isotropic resolution in-plane, 1.20 mm slice thickness (no gap), TE = 16 ms, TR = 2000 ms, α = 80°, FoV = 152 x 170 x 69 mm^3^, echo spacing = 0.78 ms, readout bandwidth = 1476 Hz/Px, GRAPPA = 3, and Partial Fourier (PF) of 6/8 in phase-encoding direction. Functional volumes (LGN localizer: 1 run of 136 volumes; M/P mapping experiment: 4 runs of 144 volumes each; motion experiment: 1 run of 130 volumes) were acquired with partial brain coverage (40 transverse slices). The number of slices and/or flip angle were adjusted in some participants (n = 4 controls and n = 5 DD) due to restrictions in energy absorption (i.e., specific absorption rate) typically associated with high-field MRI (minimum number of slices = 33, minimum flip angle = 69°). The reduction of the flip angle in these participants was well within the normal range of actual flip angle variation throughout the brain at 7 Tesla (15). In addition, we acquired one whole-brain EPI image with matching parameters to facilitate registrations between the functional and structural MRI data. To correct images for geometric distortions induced by magnetic field inhomogeneity, in each MRI session we acquired two gradient-echo datasets (Δ_TE_ = 1.02 ms) from which session-specific B0 field-maps (voxel displacement) were computed.

#### High-Resolution 7 Tesla Structural MRI Acquisition

High-resolution whole-brain structural MRI data, including a conventional T1-weighted image and a quantitative T_1_ map, were obtained using a 3D-MP2RAGE sequence (16) with the following imaging parameters: 700 μm isotropic resolution, TE = 2.45 ms, TR = 5000 ms, TI_1_/TI_2_ = 900/2750 ms, α_1_/α_2_ = 5/3°, FoV = 224 x 224 x 168 mm^3^, echo spacing = 6.8 ms, readout bandwidth = 250Hz/Px, GRAPPA = 2, and 6/8 PF in phase-encoding direction. The acquisition took 10:57 minutes.

#### Preprocessing of fMRI Data

Individual volumes of each run of the M/P mapping experiment and of the visual motion experiment were realigned to the first volume of the LGN localizer and unwarped based on the session-specific field-maps to correct for motion artifacts and EPI distortions. The whole-brain EPI was then also co-registered to this volume and subsequently used as the reference image for registering the structural to the functional data. Unwarped functional data in native space were then smoothed with a Gaussian filter with a FWHM (full width at half maximum) matching the voxel size (i.e., 1.25 x 1.25 x 1.2 mm). Times-series of each voxel were high-pass filtered (1/128 Hz cutoff) to remove low-frequency noise and signal drift. The resulting images were used for the definition of LGN M/P subdivisions and the ROI analyses (see ‘fMRI Data Analysis’ and ‘ROI Analyses’).

As part of the quality control analysis for the M/P subdivisions in native space, the unwarped functional data were also normalized into standard space (MNI, Montreal Neurological Institute). For this, the anatomical image was segmented into six tissue probability maps (gray matter, white matter, cerebrospinal fluid, soft tissue, bones, image background). These tissue class images were then non-linearly registered to the 1mm MNI brain template and derived registration parameters were applied to the functional data. The registered functional data were then smoothed with a Gaussian filter with a FWHM matching the voxel size. Finally, times-series of each voxel were high-pass filtered at a 1/128 Hz cutoff.

#### Head Motion

Head motion was assessed by computing the maximum translational and rotational displacements across each run of each fMRI experiment from the 6 motion parameters (3 translation, 3 rotation) obtained from SPM (17). Maximum translational displacements (TD) corresponded to the maximum difference between total TDs calculated as the square root of the sum of squared x, y, and z-direction displacements. Maximum rotational displacements (RD) corresponded to the maximum difference between total RDs calculated as the sum of the absolute RDs in the three directions. We also computed the mean framewise displacement (FD), which accounts for the mean translational and rotational head motion between adjacent slices. RDs were converted from degrees to millimeters assuming a spherical surface with a 50 mm radius. Independent t-tests comparing control and DD participants on these displacement measures (i.e., TD, RD, and FD) for each fMRI experiment revealed no significant group differences (LGN localizer: all *p*’s ≥ .10, M/P mapping experiment: all *p*’s ≥ .16, Motion experiment: all *p*’s ≥ .07).

#### fMRI Data Analysis

For each fMRI experiment, preprocessed data in native space were analyzed using single-participant general linear models (GLM) for block designs (18). For each participant and fMRI experiment, the two conditions of interest (i.e., LGN Localizer: left hemifield checkerboard vs. right hemifield checkerboard; M/P mapping experiment: M-stimulation vs. P-stimulation; Motion experiment: motion vs. static) were modeled as boxcar functions convolved with the canonical hemodynamic response function. For the M/P mapping experiment, data from the four runs were concatenated into a single session and additional regressors were added to account for between-run variance (17). Motion parameters (three translation and three rotation) derived from realignment and 16 physiological parameters obtained from the PhysIO toolbox (19) were also modeled as regressors of no interest to account for motion and cardio-respiratory-related variance. The physiological regressors included models of cardiac (6 regressors) and respiratory phases (8 regressors) computed using Fourier expansions of different order, based on RETROICOR (20). Physiological regressors also included models of heart rate variability (21) and respiratory volume per time (22). Including such models of physiological noise and motion parameters as nuisance regressors has been shown to substantially increase the signal-to-noise ratio in the LGN at 7 Tesla (23). Due to technical problems, physiological parameters of 6 participants could not be acquired and were not considered in the respective design matrices.

#### LGN Definition

To segment the LGN in each individual participant and to demarcate it from adjacent visual brain structures, we leveraged a publicly available, high-resolution 7T probabilistic LGN atlas (24). This atlas is available in high-resolution 0.4 mm template space as well as in MNI 1mm standard space (25). To map the bilateral LGNs in each participant’s native space, we used the command *antsRegistration* as implemented in the Advanced Normalization Tools (ANTs) software package (26), to register the high-resolution LGN atlas template to each participant’s T_1_ image. The registrations were run with rigid and affine linear registrations in combination with nonlinear symmetric normalization (SyN). All registrations were visually inspected for potential misalignments. In some participants, the local vessel architecture around the LGN affected the quality of the registrations and required the use of additional landmark information (i.e., medial-lateral and inferior-superior LGN extent in two central LGN slices) in the linear registration initialization step. After quality control, the obtained registration parameters were applied to the left and right probabilistic LGN atlases for each participant. Following registrations of these masks to the functional image data, the registered probabilistic LGN masks were set to a threshold of 35% overlap to confine final entire LGN ROI sizes to anatomically plausible volumes (left LGN: 128.2 ±17.4 mm^3^ in controls vs. 124.5 ±14.0 mm^3^ in DD; right LGN: 136.1 ±17.0 mm^3^ in controls vs. 132.4 ±17.7 mm^3^ in DD) (27). In addition, we also verified that the structurally defined entire LGN ROIs coincided with the functional LGN activations derived from the LGN localizer experiment. This was the case in all participants. These participant-specific entire LGN ROIs were used to mask the individual Beta M-P maps to subsequently define M and P subdivisions via 20/80% volume thresholding.

#### Quality Control of LGN M/P Subdivisions

We performed two main quality control analyses to assess the localization accuracy of the identified LGN M and P subdivisions:

1. To assess the structural plausibility of the M/P subdivisions, we first computed Beta M-P maps for each participant in MNI standard space. These maps were then masked with an openly available LGN probabilistic atlas (at a threshold of 35% overlap) in MNI 1mm standard space (25), and M- and P-LGN were computed via 20/80% volume thresholding. For each participant, we next calculated the centers of mass of the M- and P-LGN as a proportion of individual LGN extent. Based on prior anatomical knowledge, we expected the M-LGN to be located more medially than the P-LGN (17, 24). This was the case for the majority of participants and those who did not meet this criterion (n = 4 controls, n = 1 DD) were excluded from all further analyses. We performed this control analysis in MNI standard space to account for potential differences in LGN orientation between participants (due to differences in EPI angulation) in native space.
2. We also assessed the functional plausibility of the identified M- and P-LGN by examining their response to visual motion from the motion experiment. Based on the known response properties of LGN M and P neurons, functional responses to visual motion were expected to be stronger in the identified M- than P-LGN (28).

#### Statistical Analyses

Mean % signal change (PSC) values were submitted to mixed-design analyses of variance (ANOVA) for statistical analysis. Pairwise comparisons were performed using independent or paired t-tests, where appropriate. Effect sizes for ANOVAs and t-tests were calculated using partial eta squared 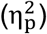 and Cohen’s d, respectively. Correlation analyses between PSC and RANln ability were performed using Pearson’s correlations. Data were checked for normality using the Shapiro-Wilk test (29). One female DD participant was excluded from the analysis of sex differences in M-LGN response in DD because her M-LGN difference score was > 2 SDs away from the group mean. For all statistical tests, the significance level α was set to 0.05, and Bonferroni-correction was applied as described in the main text.

### SI Results

#### Behavioral performance

Behavioral performance (% of correct responses) during the M/P mapping experiment (report of contrast decrements within each block) was analyzed using a mixed-design ANOVA with group (controls/DD) as between-subject factor and stimulustype (M-stimulus/P-stimulus) as within-subject factor. There was no significant main effect of group (*F*(1,41) = 0.614, *p* = .438, 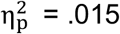), no main effect of stimulus-type (*F*(1,41) = 0.420, *p* = .520, 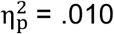) nor an interaction between both factors (*F*(1,41) = 0.006, *p* = .937, 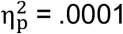). Mean performance across groups and stimuli was 43.91 ± 21.56%. This low performance could be due to errors in the use of the response keys: participants frequently reported that they found the response keys counterintuitive, tending toward not pressing any key when they detected 0 targets and using their index, middle, and ring fingers to report 1, 2, and 3 targets, respectively, thereby shifted the response keys. Such a strategy could explain the low overall accuracy. Previous M/P-mapping experiments using the exact same design reported percent correct around 71-75%, which we assume is due to running the experiment on the same participants multiple times as well as including two study authors (Denison et al., 2014).

**Table S1.**
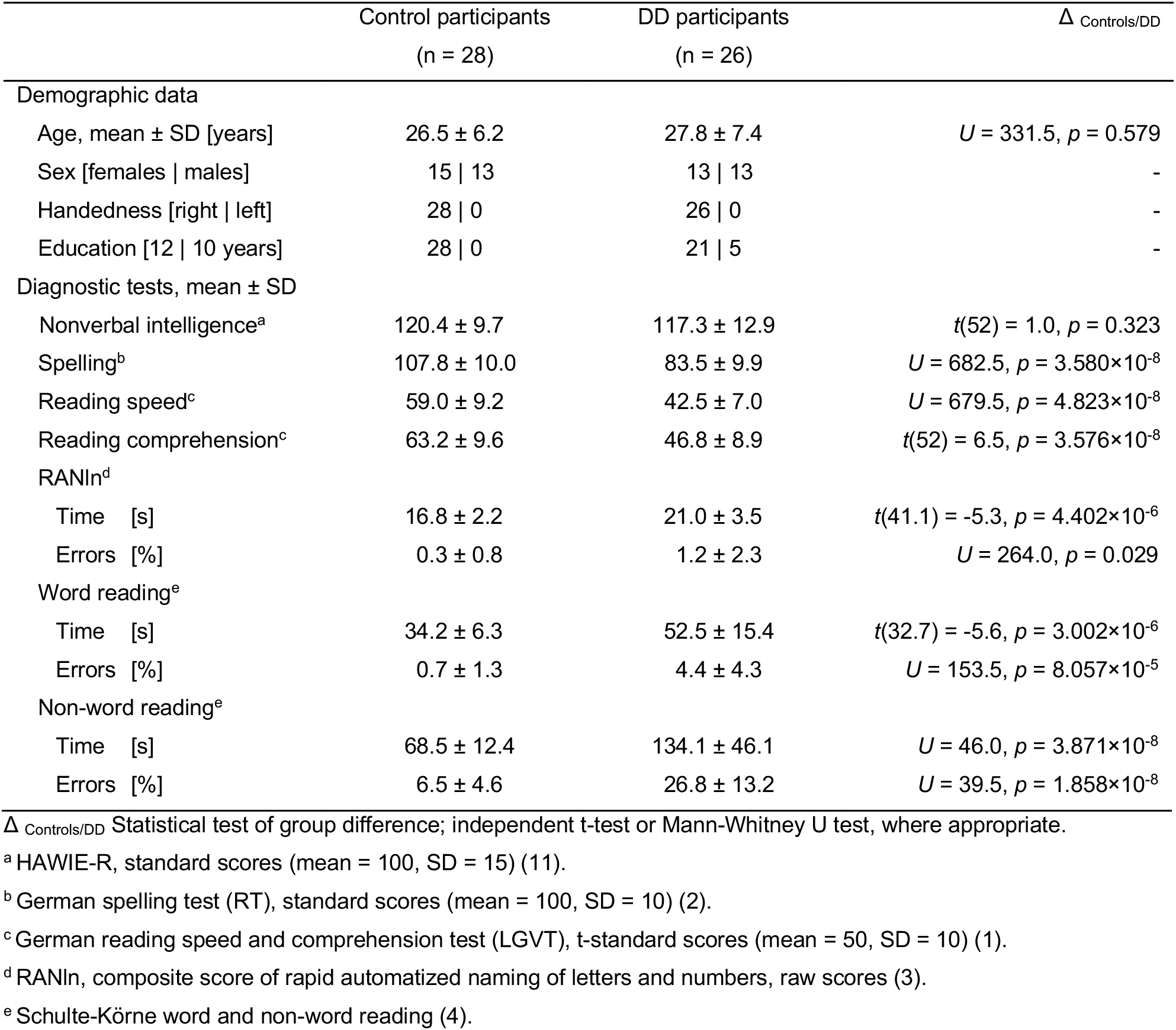
Demographic data and diagnostic test performance in controls and dyslexics.

## Notes

**Competing Interest Statement:** The authors declare no competing interests.

https://osf.io/bge75/

### Competing Interest Statement

The authors have declared no competing interest.

## References

1. B. A. Shaywitz, S. E. Shaywitz, The American experience: towards a 21st century definition of dyslexia. Oxf Rev Educ 46, 454–471 (2020).

2. S. E. Shaywitz, Dyslexia. New England Journal of Medicine 338, 307–312 (1998).

3. R. L. Peterson, B. F. Pennington, Developmental dyslexia. The Lancet 379, 1997–2007 (2012).

4. M. S. Livingstone, G. D. Rosen, F. W. Drislane, A. M. Galaburda, Physiological and anatomical evidence for a magnocellular defect in developmental dyslexia. Proceedings of the National Academy of Sciences 88, 7943–7947 (1991).

5. A. M. Galaburda, M. T. Menard, G. D. Rosen, Evidence for aberrant auditory anatomy in developmental dyslexia. Proceedings of the National Academy of Sciences 91, 8010–8013 (1994).

6. J. J. Nassi, E. M. Callaway, Parallel processing strategies of the primate visual system. Nat Rev Neurosci 10, 360–372 (2009).

7. A. E. Herman, A. M. Galaburda, R. Holly Fitch, A. R. Carter, G. D. Rosen, Cerebral microgyria, thalamic cell size and auditory temporal processing in male and female rats. Cerebral Cortex 7, 453–464 (1997).

8. G. D. Rosen, B. Mesples, M. Hendriks, A. M. Galaburda, Histometric changes and cell death in the thalamus after neonatal neocortical injury in the rat. Neuroscience 141, 875–888 (2006).

9. N. Tschentscher, A. Ruisinger, H. Blank, B. Díaz, K. von Kriegstein, Reduced structural connectivity between left auditory thalamus and the motion-sensitive planum temporale in developmental dyslexia. Journal of Neuroscience 39, 1720–1732 (2019).

10. C. Müller-Axt, A. Anwander, K. von Kriegstein, Altered structural connectivity of the left visual thalamus in developmental dyslexia. Current Biology 27, 3692–3698 (2017).

11. T. J. Andrews, S. D. Halpern, D. Purves, Correlated size variations in human visual cortex, lateral geniculate nucleus, and optic tract. Journal of Neuroscience 17, 2859–2868 (1997).

12. C. Müller-Axt, et al., Mapping the human lateral geniculate nucleus and its cytoarchitectonic subdivisions using quantitative MRI. Neuroimage 244, Article 118559 (2021).

13. F. Ramus, Developmental dyslexia: Specific phonological deficit or general sensorimotor dysfunction? Curr Opin Neurobiol 13, 212–218 (2003).

14. R. N. Denison, A. T. Vu, E. Yacoub, D. A. Feinberg, M. A. Silver, Functional mapping of the magnocellular and parvocellular subdivisions of human LGN. Neuroimage 102, 358–369 (2014).

15. E. S. Norton, M. Wolf, Rapid automatized naming (RAN) and reading fluency: Implications for understanding and treatment of reading disabilities. Annu Rev Psychol 63, 427–452 (2012).

16. B. Díaz, F. Hintz, S. J. Kiebel, K. von Kriegstein, Dysfunction of the auditory thalamus in developmental dyslexia. Proc Natl Acad Sci U S A 109, 13841–13846 (2012).

17. G. D. Rosen, A. E. Herman, A. M. Galaburda, Sex differences in the effects of early neocortical injury on neuronal size distribution of the medial geniculate nucleus in the rat are mediated by perinatal gonadal steroids. Cerebral Cortex 9, 27–34 (1999).

18. F. Ramus, Neurobiology of dyslexia: a reinterpretation of the data. Trends Neurosci 27, 720–726 (2004).

19. M. B. Denckla, R. G. Rudel, Rapid “automatized” naming (R.A.N.): Dyslexia differentiated from other learning disabilities. Neuropsychologia 14, 471–479 (1976).

20. S. Rima, G. Kerbyson, E. Jones, M. C. Schmid, Advantage of detecting visual events in the right hemifield is affected by reading skill. Vision Res 169, 41–48 (2020).

21. T. M. Evans, D. L. Flowers, E. M. Napoliello, G. F. Eden, Sex-specific gray matter volume differences in females with developmental dyslexia. Brain Struct Funct 219, 1041–1054 (2014).

22. A. J. Krafnick, E. M. Napoliello, D. L. Flowers, G. F. Eden, The Role of Brain Activity in Characterizing Successful Reading Intervention in Children With Dyslexia. Front Neurosci 16 (2022).

23. A. J. Krafnick, T. M. Evans, Neurobiological Sex Differences in Developmental Dyslexia. Front Psychol 9 (2019).

24. M. Corbetta, et al., A Common Network of Functional Areas for Attention and Eye Movements. Neuron 21, 761–773 (1998).

25. M. T. de Schotten, et al., A lateralized brain network for visuospatial attention. Nat Neurosci 14, 1245–1246 (2011).

26. R. T. Born, D. C. Bradley, Structure and function of visual area MT. Annu Rev Neurosci 28, 157–189 (2005).

27. R. Hari, Left minineglect in dyslexic adults. Brain 124, 1373–1380 (2001).

28. R. Hari, H. Renvall, Impaired processing of rapid stimulus sequences in dyslexia. Trends Cogn Sci 5, 525–532 (2001).

29. I. Heth, M. Lavidor, Improved reading measures in adults with dyslexia following transcranial direct current stimulation treatment. Neuropsychologia 70, 107–113 (2015).

30. A. Battisti, et al., Effects of a short and intensive transcranial direct current stimulation treatment in children and adolescents with developmental dyslexia: A crossover clinical trial. Front Psychol 13 (2022).

31. C. Hutton, et al., The impact of physiological noise correction on fMRI at 7 T. Neuroimage 57, 101–112 (2011).

32. K. M. O’Craven, B. R. Rosen, K. K. Kwong, A. Treisman, R. L. Savoy, Voluntary attention modulates fMRI activity in human MT–MST. Neuron 18, 591–598 (1997).

33. B. B. Avants, C. L. Epstein, M. Grossman, J. C. Gee, Symmetric diffeomorphic image registration with cross-correlation: Evaluating automated labeling of elderly and neurodegenerative brain. Med Image Anal 12, 26–41 (2008).

34. C. R. Pernet, Misconceptions in the use of the General Linear Model applied to functional MRI: a tutorial for junior neuro-imagers. Front Neurosci 8 (2014).

## SI References

1. W. Schneider, M. Schlagmüller, M. Ennemoser, LGVT 6-12: Lesegeschwindigkeits-und - verständnistest für die Klassen 6–12 (Hogrefe, 2007).

2. M. Kersting, K. Althoff, Rechtschreibungstest: RT (Hogrefe, 2004).

3. M. B. Denckla, R. G. Rudel, Rapid ‘automatized’ naming (R.A.N.): Dyslexia differentiated from other learning disabilities. Neuropsychologia 14, 471–479 (1976).

4. G. Schulte-Körne, *Lese-Rechtschreibstörung und Sprachwahrnehmung*. Psychometrische und neurophysiologische Untersuchungen zur Legasthenie. (Waxmann, 2001).

5. S. Baron-Cohen, S. Wheelwright, R. Skinner, J. Martin, E. Clubley, The Autism-Spectrum Quotient (AQ): Evidence from Asperger syndrome/high-functioning autism, males and females, scientists and mathematicians. J Autism Dev Disord 31, 5–17 (2001).

6. B. Butterworth, “Developmental Dyscalculia” in Handbook of Mathematical Cognition, J. I. D. Campbell, Ed. (Psychology Press, 2005), pp. 455–468.

7. R. S. Shalev, Developmental Dyscalculia. J Child Neurol 19, 765–771 (2004).

8. M. Bach, The Freiburg Visual Acuity Test - Automatic measurement of visual acuity. Optometry and Vision Science 73, 49–53 (1996).

9. M. Bach, The Freiburg Visual Acuity Test - Variability unchanged by post-hoc re-analysis. Graefe’s Archive for Clinical and Experimental Ophthalmology 245, 965–971 (2006).

10. A. Colenbrander, “Visual standards: aspects and ranges of vision loss with emphasis on population surveys. Report for the International Council of Ophthalmology.” (2002).

11. U. Tewes, Hamburg-Wechsler-Intelligenztest für Erwachsene (HAWIE-R), rev 1991 (Hans Huber, 1991).

12. D. G. Pelli, The VideoToolbox software for visual psychophysics: transforming numbers into movies. Spat Vis 10, 437–442 (1997).

13. D. H. Brainard, The Psychophysics Toolbox. Spat Vis 10, 433–436 (1997).

14. J. Eaton, D. Bateman, S. Hauberg, R. Wehbring, GNU Octave Version 4.2.0 Manual: A high-level interactive language for numerical computations. Available online at: http://www.gnu.org/software/octave/doc/interpreter/ (2016).

15. J. T. Vaughan, et al., 7T vs. 4T: RF power, homogeneity, and signal-to-noise comparison in head images. Magn Reson Med 46, 24–30 (2001).

16. J. P. Marques, et al., MP2RAGE, a self bias-field corrected sequence for improved segmentation and T1-mapping at high field. Neuroimage 49, 1271–1281 (2010).

17. R. N. Denison, A. T. Vu, E. Yacoub, D. A. Feinberg, M. A. Silver, Functional mapping of the magnocellular and parvocellular subdivisions of human LGN. Neuroimage 102, 358–369 (2014).

18. K. J. Friston, et al., Statistical parametric maps in functional imaging: A general linear approach. Hum Brain Mapp 2, 189–210 (1994).

19. L. Kasper, et al., The PhysIO toolbox for modeling physiological noise in fMRI data. J Neurosci Methods 276, 56–72 (2017).

20. G. H. Glover, T.-Q. Li, D. Ress, Image-based method for retrospective correction of physiological motion effects in fMRI: RETROICOR. Magn Reson Med 44, 162–167 (2000).

21. C. Chang, J. P. Cunningham, G. H. Glover, Influence of heart rate on the BOLD signal: The cardiac response function. Neuroimage 44, 857–869 (2009).

22. R. M. Birn, M. A. Smith, T. B. Jones, P. A. Bandettini, The respiration response function: The temporal dynamics of fMRI signal fluctuations related to changes in respiration. Neuroimage 40, 644–654 (2008).

23. C. Hutton, et al., The impact of physiological noise correction on fMRI at 7 T. Neuroimage 57, 101–112 (2011).

24. C. Müller-Axt, et al., Mapping the human lateral geniculate nucleus and its cytoarchitectonic subdivisions using quantitative MRI. Neuroimage 244, Article 118559 (2021).

25. C. Müller-Axt, et al., High-Resolution LGN Atlas. Open Science Framework Repository. doi:10.17605/OSF.IO/TQAYF (2020).

26. B. B. Avants, C. L. Epstein, M. Grossman, J. C. Gee, Symmetric diffeomorphic image registration with cross-correlation: Evaluating automated labeling of elderly and neurodegenerative brain. Med Image Anal 12, 26–41 (2008).

27. T. J. Andrews, S. D. Halpern, D. Purves, Correlated size variations in human visual cortex, lateral geniculate nucleus, and optic tract. Journal of Neuroscience 17, 2859–2868 (1997).

28. J. J. Nassi, E. M. Callaway, Parallel processing strategies of the primate visual system. Nat Rev Neurosci 10, 360–372 (2009).

29. P. Royston, Approximating the Shapiro-Wilk W-test for non-normality. Stat Comput 2, 117–119 (1992).

